# IL-1R1 Blockade Boosts CD40 Agonist Immune Responses but Fails to Improve Efficacy or Reduce Hepatotoxicity in Pancreatic Cancer

**DOI:** 10.1101/2025.02.23.639774

**Authors:** Akash Boda, Irfan N. Bandey, Saikat Chowdhury, Sadhna Aggarwal, Mekala Venugopala, Natalie Wall Fowlkes, Jason Roszik, Michael A. Curran, Van Karlyle Morris, Scott Kopetz, Manisha Singh

**Affiliations:** Department of Immunology, The University of Texas MD Anderson Cancer Center, Houston, Texas, USA; Department of Gastrointestinal Medical Oncology, The University of Texas MD Anderson Cancer Center, Houston, Texas, USA; Department of Thoracic Radiation Oncology, The University of Texas MD Anderson Cancer Center, Houston, Texas, USA; Institute for Clinical and Translational Research, Dan L. Duncan Comprehensive Cancer Center, Baylor College of Medicine, Houston, Texas, USA; Department of Veterinary Medicine and Surgery, The University of Texas MD Anderson Cancer Center, Houston, Texas, USA; Department of Melanoma, The University of Texas MD Anderson Cancer Center, Houston, Texas, USA

## Abstract

Pancreatic ductal adenocarcinoma (PDAC) has a poor survival rate and limited treatments. Agonistic CD40 antibodies are promising, but clinical trials have shown only modest efficacy and significant hepatotoxicity. We previously reported that IL-1 pathway blockade enhances agonistic CD40 antibody efficacy against melanoma by depleting polymorphonuclear myeloid-derived suppressor cells (PMN-MDSCs; CD11b^+^Ly6C^+^Ly6G^+^). Because PMN-MDSCs also cause liver toxicity, we investigated the impact of IL-1R1 blockade on the efficacy and toxicity of agonistic CD40 antibody therapy in PDAC. Agonistic CD40 antibody therapy induced immune activation and significantly prolonged survival in orthotopic PDAC-bearing mice. IL-1R1 blockade monotherapy downregulated innate and adaptive immune response and exacerbated tumor growth. Although combination therapy upregulated several immune-related pathways and boosted innate and adaptive immune responses. IL-1R1 blockade failed to improve the overall antitumor efficacy of agonistic CD40 antibody therapy and exacerbated liver toxicity. Ly6G^+^ cell depletion in mice reduced the efficacy of agonistic CD40 antibody therapy, suggesting that Ly6G⁺ immune cells (PMN-MDSCs or neutrophils) exhibit an antitumor rather than immunosuppressive role in PDAC. Our findings underscore the complex role of IL-1 signaling in modulating immune responses in PDAC and caution against pursuing IL-1R1 blockade, either as monotherapy or combined with agonistic CD40 antibodies, in clinical trials for PDAC.

## Introduction

Pancreatic ductal adenocarcinoma (PDAC) is incurable and has a 5-year survival rate of only 10% in the United States ^1^. PDAC is predicted to become the second leading cause of cancer-related death by 2030 ^2^. The primary cause of the high lethality of PDAC is resistance to therapy. Immunotherapy, including immune checkpoint blockade, such as treatment with an anti-PD-1 antibody, is effective against many solid tumors, but PDAC does not respond to immune checkpoint blockade ^3, 4^.

PDAC is characterized by extensive accumulation of myeloid cells. Agonistic CD40 antibody therapy activates PDAC-associated myeloid cells through immune co-stimulatory receptor CD40, inducing both innate and tumor-specific immunity and sensitizing the tumor microenvironment to other immunotherapies ^5, 6^. Therefore, the agonistic CD40 antibody is considered the most promising immunotherapy for PDAC. However, clinical trials of several agonistic CD40 antibodies with or without immune checkpoint blockade showed only moderate therapeutic efficacy, and most patients had relapse after an initial response ^7–9^. The combination of agonistic CD40 antibody therapy and immune checkpoint blockade induces dose-limiting immune-related adverse events, such as cytokine storm and liver damage ^10–13^. To enhance the antitumor activity and lower the toxicity of agonistic CD40 antibody therapy, it is crucial to discover and target the molecules and pathways that cause resistance to such therapy and therapy-mediated toxicity.

Inflammation is associated with cancer initiation and progression ^14^; however, the underlying mechanism is not clear. In addition, the role of inflammation in resistance to immunotherapies has not been well studied. IL-1 is a major pro-inflammatory cytokine present in the inflammatory tumor microenvironment ^15^. IL-1 increases tumor invasiveness and metastasis. The two forms of IL-1, IL-1α and IL-1β, are derived from different genes but are functionally similar, and both bind to IL-1 receptor type 1 (IL-1R1) ^16, 17^. IL-1 induces accumulation of myeloid-derived suppressor cells (MDSCs) ^18^ and regulatory T cells ^19^ at the site of inflammation, but the importance of this mechanism in the tumor microenvironment is not well studied. IL-1 induces the production of IL-6, IL-8, IL-17, and C-reactive protein ^20–22^, which are associated with immune-related adverse events and development of resistance to immunotherapies ^23–27^.

We previously reported that agonistic CD40 antibody therapy induced IL-1α in the melanoma microenvironment, which conferred adaptive resistance to agonistic CD40 antibody therapy through polymorphonuclear MDSCs (PMN-MDSCs) ^28^. Although phenotypically similar to activated neutrophils, PMN-MDSCs are pathologically activated and exhibit immunosuppressive properties. In the tumor microenvironment, activated neutrophils contribute to antitumor immunity, whereas immunosuppressive PMN-MDSCs promote tumor growth^29^.

Agonistic CD40 antibody therapy induces liver damage through tumor-derived inflammatory MDSCs and PMN-MDSCs ^12, 30^. We previously reported that inflammatory monocytes were the key cells that produced IL-1 in response to agonistic CD40 antibody and IL-1–derived PMN-MDSCs in the tumor ^28^. The aim of the current study was to determine the effect of IL-1R1 blockade on the efficacy and toxicity of agonistic CD40 antibody therapy in PDAC.

## Results

### Agonistic CD40 antibody therapy reshapes the tumor microenvironment and improves survival in an orthotopic PDAC mouse model

To assess the therapeutic potential of agonistic CD40 antibody therapy in PDAC, we first evaluated its activity and efficacy in an orthotopic PDAC mouse model. Bulk transcriptomic analysis revealed a significantly higher number of differentially expressed genes in the agonistic CD40 antibody therapy group compared with the isotype control group, with 7,362 genes showing altered expression (Fig. 1a).

**Figure 1.**
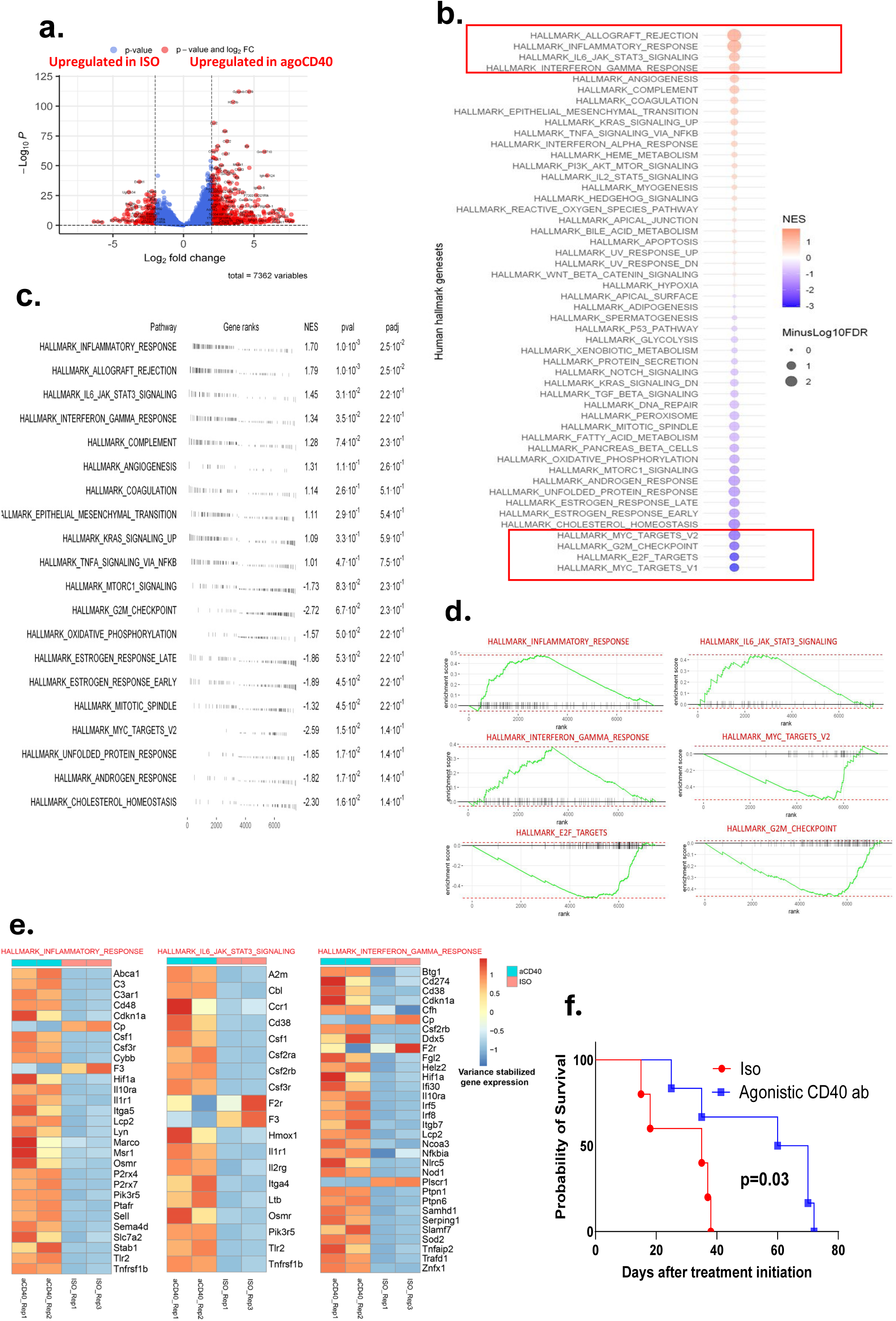
Agonistic CD40 antibody therapy induces immune activation and prolongs survival in pancreatic ductal adenocarcinoma (PDAC)-bearing mice. Mice bearing 9-day-old orthotopic PDAC tumors (confirmed by luminescence imaging) were treated every 3 days with agonistic CD40 antibody or isotype control (200 µg/mouse, intraperitoneal injection). Bulk transcriptomic analysis of tumors was performed 2 weeks after treatment initiation (n = 2 mice/group). (a) Volcano plot highlighting differentially expressed genes between CD40-treated and isotype control tumors. (b) Gene set enrichment analysis of bulk RNA sequencing data showing hallmark pathways enriched in CD40-treated tumors compared with controls. Upregulated pathways included inflammatory response, IFN-γ response, and IL-6/JAK-STAT signaling, and oncogenic pathways such as MYC targets V2, E2F targets, and G2M checkpoint were significantly downregulated (false discovery rate [FDR] < 0.05). (c) Summary of gene ranks, normalized enrichment scores (NES), p values, and adjusted p values (padj) for hallmark pathways. (d) Enrichment plots for select hallmark pathways, illustrating the distribution of pathway-associated genes in the ranked gene list. (e) Heatmap of hallmark gene signatures, depicting relative expression across samples, normalized by row Z-scores. (f) Kaplan-Meier survival curves of orthotopic PDAC-bearing mice treated with agonistic CD40 antibody or isotype control (log-rank test, n = 5-6 mice/group), Data is representative of at least 2 independent experiments.

Pathway enrichment analysis highlighted a profound immunomodulatory effect of agonistic CD40 antibody therapy. Key immunologic hallmark pathways, including the inflammatory response, IFN-γ response, and IL-6/JAK-STAT signaling pathways, were significantly upregulated (p < 0.01), suggesting a robust activation of both innate and adaptive immunity. In contrast, oncogenic pathways such as MYC targets V2, E2F targets, and the G2M checkpoint were markedly downregulated (p < 0.01), indicating potential tumor-suppressive effects at the transcriptional level (Fig. 1b-c). Gene set enrichment analysis further confirmed these findings (Fig. 1d), and genes associated with these immune-related pathways are detailed in Fig. 1e.

Agonistic CD40 antibody therapy led to a significant extension in survival of orthotopic PDAC-bearing mice compared with isotype antibody-treated controls (Fig. 1f). This suggests that agonistic CD40 antibody therapy not only remodels the tumor microenvironment by promoting pro-inflammatory and antitumor immune responses but also effectively suppresses oncogenic signaling, ultimately inhibiting tumor progression and improving survival outcomes.

### IL-1R1 blockade enhances agonistic CD40 antibody–induced immunity but fails to improve antitumor efficacy in PDAC

To assess the impact of IL-1R1 blockade on the efficacy of agonistic CD40 antibody therapy, we first examined the expression of IL-1R1 and IL-1α/β in orthotopic PDAC tumors. These tumors expressed IL-1R1 and IL-1α in response to agonistic CD40 antibody (Fig. 2a). Next, we evaluated the antitumor efficacy of anti-IL-1R1 antibody, either as monotherapy or in combination with an agonistic CD40 antibody, in both orthotopic and subcutaneous PDAC models. We found that anti-IL-1R1 antibody monotherapy reduced survival in orthotopic PDAC-bearing mice compared with the isotype control, but treatment with the combination of anti-IL-1R1 and agonistic CD40 antibodies did not improve outcomes over agonistic CD40 antibody therapy alone (Fig. 2b). Similarly, in subcutaneous PDAC models, mice treated with anti-IL-1R1 antibodies showed accelerated tumor growth compared with the untreated group, and treatment with the combination of anti-IL-1R1 and agonistic CD40 antibodies performed worse than agonistic CD40 antibody monotherapy (Fig. 2c). Histological analysis revealed significant more necrosis in the agonistic CD40 antibody treatment group than in the combination treatment group (Fig. 2d-e).

**Figure 2.**
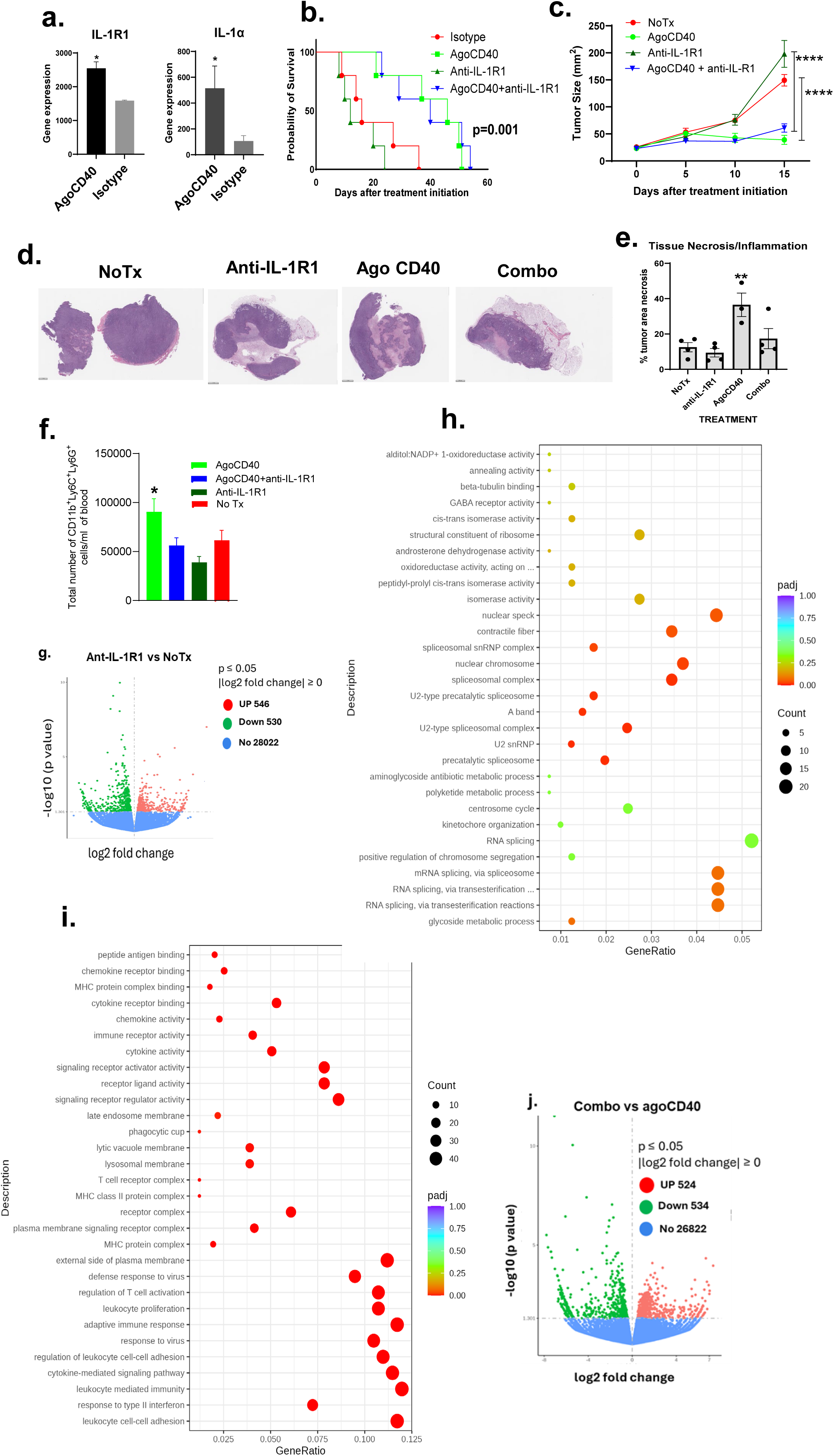
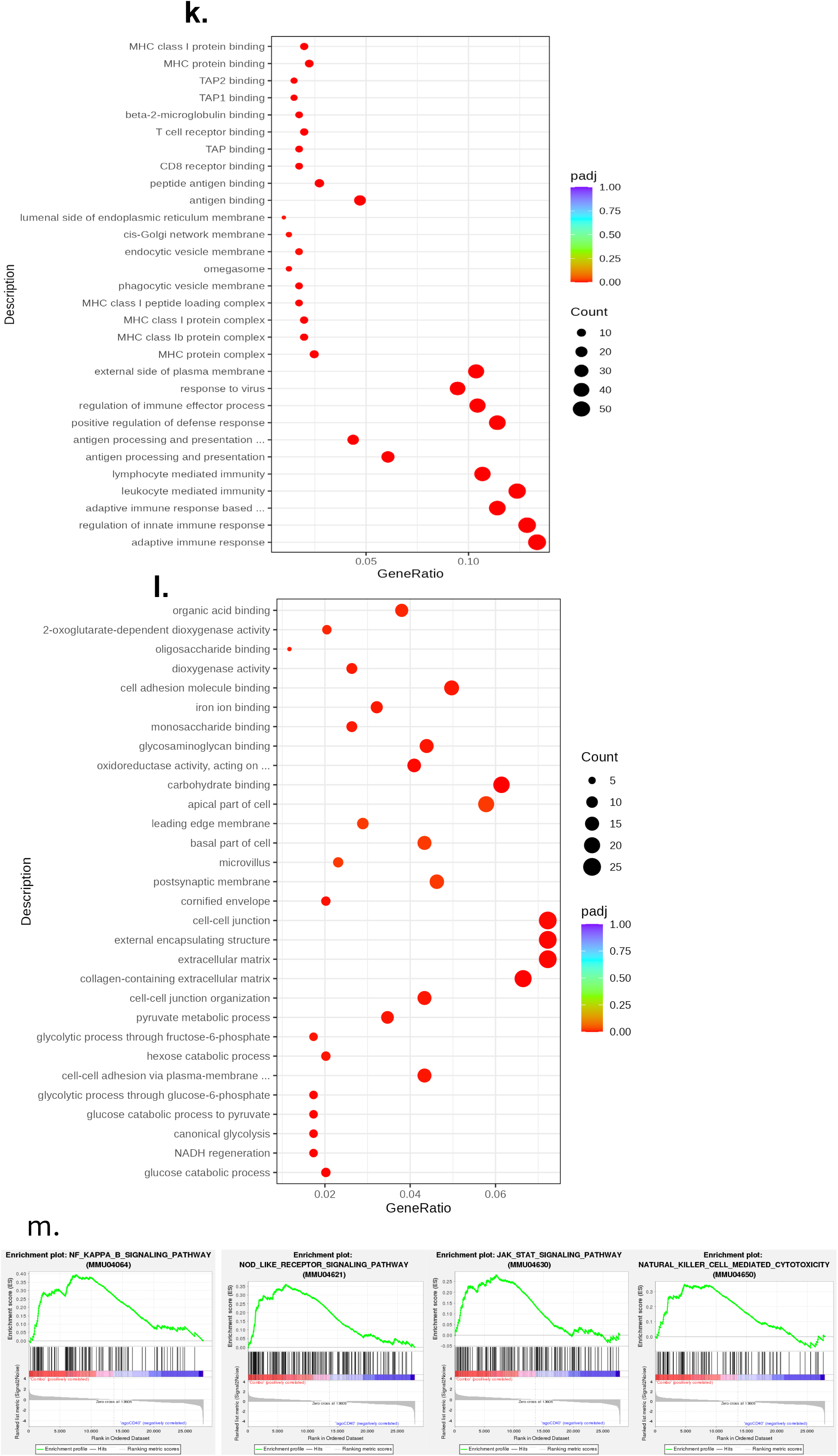
Combination therapy with agonistic CD40 and anti-IL-1R1 antibodies enhances immune activation but fails to improve therapeutic efficacy in pancreatic ductal adenocarcinoma (PDAC) compared with agonistic CD40 antibody monotherapy. Mice bearing 9-day-old orthotopic PDAC tumors (confirmed by luminescence imaging) were treated every 3 days with the indicated antibodies (200 µg/mouse, intraperitoneal). (a) Bulk transcriptomic analysis of orthotopic PDAC tumors 2 weeks after treatment initiation (mean ± SEM, n = 2 mice/group; *p < 0.05, unpaired *t* test). (b) Kaplan-Meier survival curves (log-rank test, n = 5-6 mice/group). Data is representative of at least 2 independent experiments. (c) Tumor growth kinetics in subcutaneous PDAC-bearing mice treated every 3 days with agonistic CD40 antibody, anti-IL-1R1 antibody, their combination, or left untreated (mean ± SEM, n = 5-8 mice/group; ***p < 0.0001, two-way ANOVA). (d) Representative H&E-stained tumor sections showing necrosis after 2 weeks of treatment (n = 3-4 mice/group). (e) Quantification of necrotic areas (mean ± SEM, n = 3-4 mice/group; **p < 0.01, one-way ANOVA). (f) Flow cytometry analysis of polymorphonuclear myeloid-derived suppressor (CD11b^+^Ly6C^+^Ly6G^+^) cells in peripheral blood after 2 weeks of treatment (mean ± SEM, n = 5 mice/group; *p < 0.05, unpaired Student *t* test). (g) Volcano plot of differentially expressed genes (DEGs) in subcutaneous PDAC tumors treated with anti-IL-1R1 antibody versus untreated controls (red: upregulated; green: downregulated; 546 up, 530 down; p ≤ 0.05, log2 fold change ≥ 0). (h–i) Gene ontology enrichment analysis of DEGs from (g), showing significantly upregulated (h) and downregulated (i) biological processes (padj: adjusted p value). (j) Volcano plot of DEGs comparing combination therapy (agonistic CD40 and anti-IL-1R1 antibodies) versus CD40 monotherapy (524 up, 534 down; p ≤ 0.05, log2 fold change ≥ 0). (k–l) Gene ontology enrichment analysis of DEGs from (j), highlighting upregulated (k) and downregulated (l) pathways. (m) Gene set enrichment analysis plots showing enrichment of indicated gene sets in the combination therapy group compared with agonistic CD40 antibody monotherapy.

Previously, we reported that IL-1R1 blockade reduces the number of PMN-MDSCs in melanoma ^28^. To determine whether this effect extends to PDAC, we analyzed PMN-MDSC levels in PDAC-bearing mice. Indeed, treatment with an anti-IL-1R1 antibody significantly reduced the number of PMN-MDSCs in the blood of PDAC-bearing mice treated with agonistic CD40 antibodies (Fig. 2f).

Next, we examined the impact of IL-1R1 blockade on the tumor microenvironment by comparing the bulk transcriptomes of PDAC tumors. Specifically, we analyzed tumors treated with an anti-IL-1R1 antibody versus untreated controls, as well as tumors receiving a combination of agonistic CD40 and anti-IL-1R1 antibodies versus agonistic CD40 antibody monotherapy. Through gene ontology enrichment analysis of differentially expressed genes (p ≤ 0.05; Fig. 2g), the anti-IL-1R1 antibody group showed upregulation of genes related to the spliceosome, spliceosome complex, and mRNA splicing compared with the no-treatment group (adjusted p = 0.00-0.1; Fig. 2h). In contrast, we identified several immune-related pathways and molecules that were significantly downregulated in the anti-IL-1R1 antibody group compared with the no-treatment group. These included response to type II IFN, leukocyte-mediated immunity, cytokine-mediated signaling pathways, adaptive immune response, and regulation of T-cell activation (adjusted p = 0.00; Fig. 2i).

When comparing combination therapy (agonistic CD40 plus anti-IL-1R1 antibodies) with agonistic CD40 antibody monotherapy using gene ontology enrichment analysis of differentially expressed genes (p ≤ 0.05; Fig. 2j), we found that the combination therapy group was enriched for activated innate and adaptive immune responses. This included processes such as antigen processing and presentation, leukocyte-mediated immunity, lymphocyte-mediated immunity, adaptive immune responses, and regulation of immune effector processes (adjusted p = 0.00; Fig. 2k). In contrast, many metabolic pathways were downregulated in the combination therapy group (adjusted p = 0.00; Fig. 2l).

Furthermore, gene set enrichment analysis confirmed the enrichment of genes involved in NOD-like receptor signaling, nuclear factor kappa B (NF-κB) signaling, NK cell–mediated cytotoxicity, and the JAK-STAT signaling pathway in the combination therapy group compared with the agonistic CD40 antibody monotherapy group (Fig. 2m).

Taken together, these findings suggest that although combination therapy with agonistic CD40 and anti-IL-1R1 antibodies elicits a more robust immune response than agonistic CD40 antibody monotherapy, it does not enhance the antitumor efficacy of the agonistic CD40 antibody.

### IL-1R1 blockade exacerbates agonistic CD40 antibody–induced hepatotoxicity

Agonistic CD40 antibody induces liver damage through tumor-derived PMN-MDSCs ^12, 30^. We found that blocking the IL-1 pathway reduced the number of agonistic CD40 antibody–induced PMN-MDSCs in the blood (Fig. 2e). Based on this observation, we hypothesized that inhibiting the IL-1 pathway using an anti-IL-1R1 antibody might mitigate agonistic CD40 antibody–induced hepatotoxicity.

To investigate whether agonistic CD40 antibody–induced hepatotoxicity is mediated by IL-1, we measured serum levels of aspartate aminotransferase (AST) and alanine aminotransferase (ALT) markers of liver injury in PDAC-bearing mice treated with either agonistic CD40 antibody monotherapy or the combination of agonistic CD40 and anti-IL-1R1 antibodies. Measurements were taken 48 hours, 9 days, and 3 weeks after treatment initiation.

At 48 hours, both AST and ALT levels were significantly elevated in mice treated with agonistic CD40 antibody compared with untreated controls. Unexpectedly, the addition of anti-IL-1R1 antibody to agonistic CD40 antibody therapy further exacerbated AST and ALT elevations compared with agonistic CD40 antibody monotherapy (Fig. 3a-b). Histological analysis (H&E staining) of liver sections at 48 hours further supported this finding, revealing increased immune infiltration in both the agonistic CD40 antibody monotherapy group (vs untreated controls) and combination therapy group (vs agonistic CD40 antibody monotherapy; Fig. 3c-d). These results indicate that IL-1 pathway blockade worsens agonistic CD40 antibody– induced hepatotoxicity.

**Figure 3.**
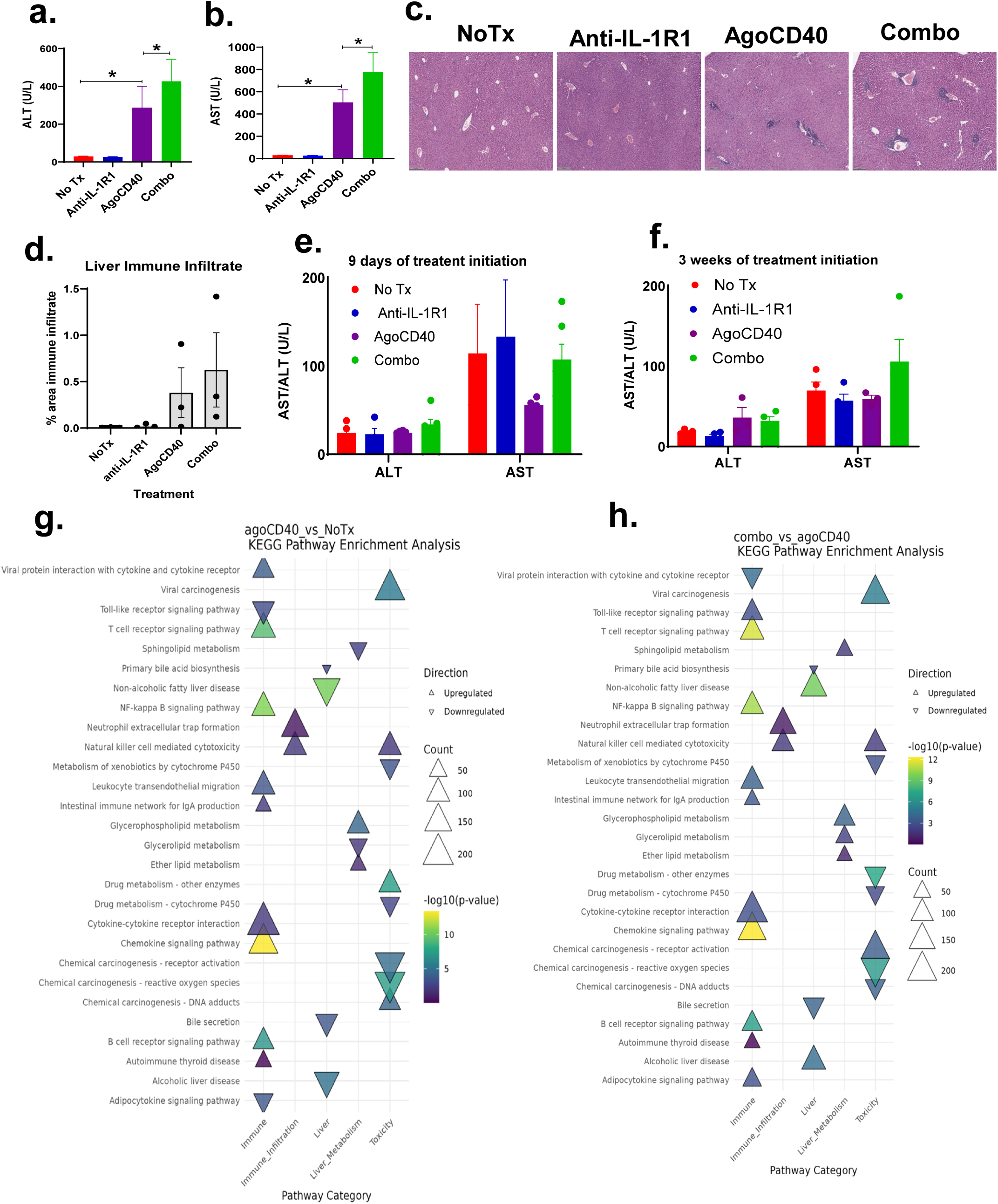
IL-1R1 blockade exacerbates agonistic CD40 antibody–induced hepatoxicity. Mice bearing 9-day-old subcutaneous pancreatic ductal adenocarcinoma (PDAC) tumors were treated as indicated (Combo: agonistic CD40 and anti-IL-1R1 antibodies). (a-b) Serum alanine aminotransferase (ALT; a) and aspartate aminotransferase (AST; b) levels were measured 48 hours after treatment initiation using the COBAS INTEGRA 400 Plus analyzer (mean ± SEM, n = 6-9 mice/group; *p < 0.01, Wilcoxon nonparametric test). Data represent results combined from two independent experiments. (c) Representative H&E-stained liver histologic images from mice, collected 1 week after treatment initiation (n = 3 mice/group, two treatments administered). (d) Cumulative quantification of immune infiltration and histologic changes (mean ± SEM, n = 3 mice/group). (e-f) Serum ALT and AST levels measured after 9 days (e) and 3 weeks (f) of treatment. (g-h) Transcriptomic profiling of the liver from PDAC-bearing mice 1 week after treatment initiation (two treatments total). KEGG pathway enrichment analysis was used to compare agonistic CD40 antibody vs no treatment (g) and combination treatment vs agonistic CD40 antibody monotherapy (h). The dot plot highlights pathways associated with liver function, toxicity, and immune infiltration. Dot size represents the number of genes involved (Count), and color intensity reflects statistical significance (-log10 p value). Upregulated pathways are marked with upward triangles (▴) and downregulated pathways with downward triangles (▾). Pathways with p < 0.05 are considered significantly enriched.

After 9 days and 3 weeks of treatment (with dosing every 3 days), AST and ALT levels were no longer elevated in either the monotherapy or combination therapy groups (Fig. 3e-f). This suggests that agonistic CD40 antibody–induced hepatotoxicity is transient and resolves over time.

To investigate the mechanisms underlying hepatotoxicity induced by agonistic CD40 antibody monotherapy and its combination with anti-IL-1R1 antibody therapy, we performed transcriptomic profiling of liver tissues. To interpret the biological significance of differentially expressed genes, we conducted KEGG pathway enrichment analysis. Our analysis identified genes and pathways associated with liver function, toxicity, and immune infiltration in both monotherapy and combination therapy groups. When comparing agonistic CD40 antibody therapy with the untreated control, we observed significant upregulation of pathways related to B-cell receptor signaling, chemokine receptor signaling, NK cell–mediated cytotoxicity, neutrophil extracellular trap formation, NF-κB signaling, glycerophospholipid metabolism, and viral carcinogenesis (Fig. 3g).

In contrast, the comparison between combination therapy (agonistic CD40 and anti-IL-1R1 antibodies) and agonistic CD40 antibody monotherapy revealed enrichment of similar immune-related pathways, including B-cell receptor signaling, chemokine receptor signaling, NK cell–mediated cytotoxicity, neutrophil extracellular trap formation, NF-κB signaling, and viral carcinogenesis. However, unlike the monotherapy group, the combination therapy group also exhibited significant upregulation of pathways related to alcoholic liver disease, non-alcoholic fatty liver disease, and chemical carcinogenesis-receptor activation (Fig. 3h), suggesting an increased risk of liver toxicity with combination therapy.

### Ly6G⁺ cells (PMN-MDSCs/neutrophils) exhibit antitumor effects in PDAC

Although agonistic CD40 antibody enhances antitumor immunity, it also induces the expansion of PMN-MDSCs, which are typically associated with immunosuppressive effects on antitumor responses. To counteract this, we investigated the impact of anti-IL-1R1 antibody therapy, which effectively reduced agonistic CD40 antibody–induced PMN-MDSCs (CD11b⁺Ly6C⁺Ly6G⁺ cells; Fig. 2e). However, despite this reduction, anti-IL-1R1 antibody therapy failed to enhance the efficacy of the agonistic CD40 antibody therapy (Fig. 2b-c).

To further clarify whether Ly6G⁺ cells play a pro-tumorigenic or anti-tumorigenic role in PDAC, we selectively depleted Ly6G⁺ cells and assessed their impact on tumor growth. Unexpectedly, Ly6G⁺ cell depletion accelerated tumor progression (Fig. 4). These findings challenge the traditional view of Ly6G⁺ cells as purely immunosuppressive and suggest that, in the context of PDAC, Ly6G⁺ cells may have a protective, antitumor immune function.

**Figure 4.**
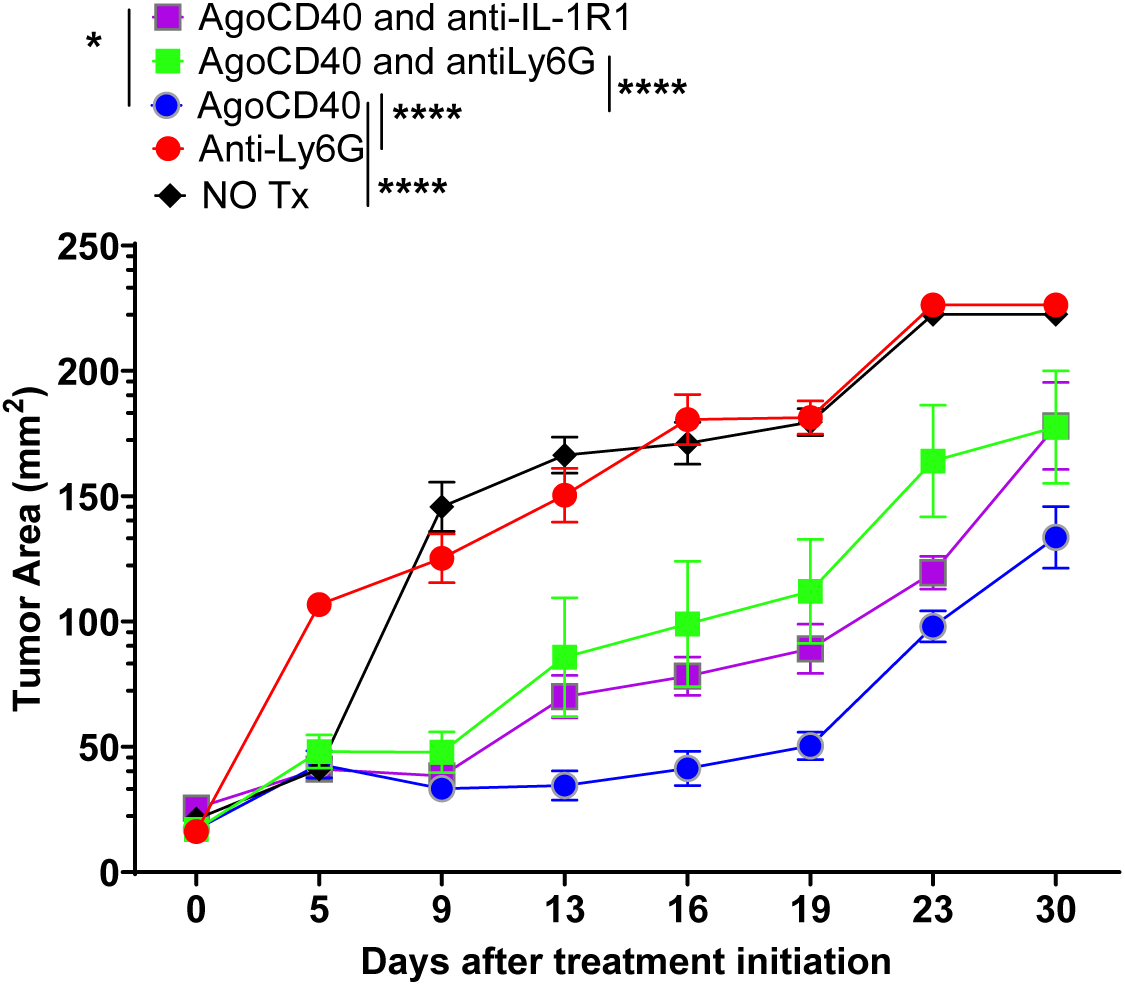
Ly6G⁺ cells (polymorphonuclear myeloid-derived suppressor cells or neutrophils) exhibit antitumor effects in pancreatic ductal adenocarcinoma (PDAC). Tumor growth kinetics are shown in mice bearing 9-day-old subcutaneous PDAC tumors treated as indicated. Tumor area (mm²) was measured over 30 days following treatment initiation. Treatment groups included no treatment (NO Tx), anti-Ly6G, agonistic CD40 antibody (AgoCD40), AgoCD40 combined with anti-Ly6G, and AgoCD40 combined with anti-IL-1R1 antibodies. Data represent mean ± SEM (n = 6-9 mice per group). Statistical significance was determined by two-way ANOVA with the Tukey post hoc test. ***p < 0.001, ****p < 0.0001.

### IL-1R1, IL-1α, and IL-1β expression are not associated with survival in human PDAC

To enhance the translational relevance of our preclinical findings, we examined the expression of IL-1R1, IL-1α, and IL-1β in human PDAC. Our analysis revealed that expression levels were significantly higher in PDAC tumors compared with normal pancreatic tissue (Fig. 5a), suggesting a potential role for IL-1 signaling in PDAC. However, survival analysis indicated that IL-1R1, IL-1α, and IL-1β expression levels did not significantly impact overall survival in PDAC patients (Fig. 5b). These findings suggest that targeting the IL-1 pathway may have limited therapeutic benefit for PDAC patients.

**Figure 5.**
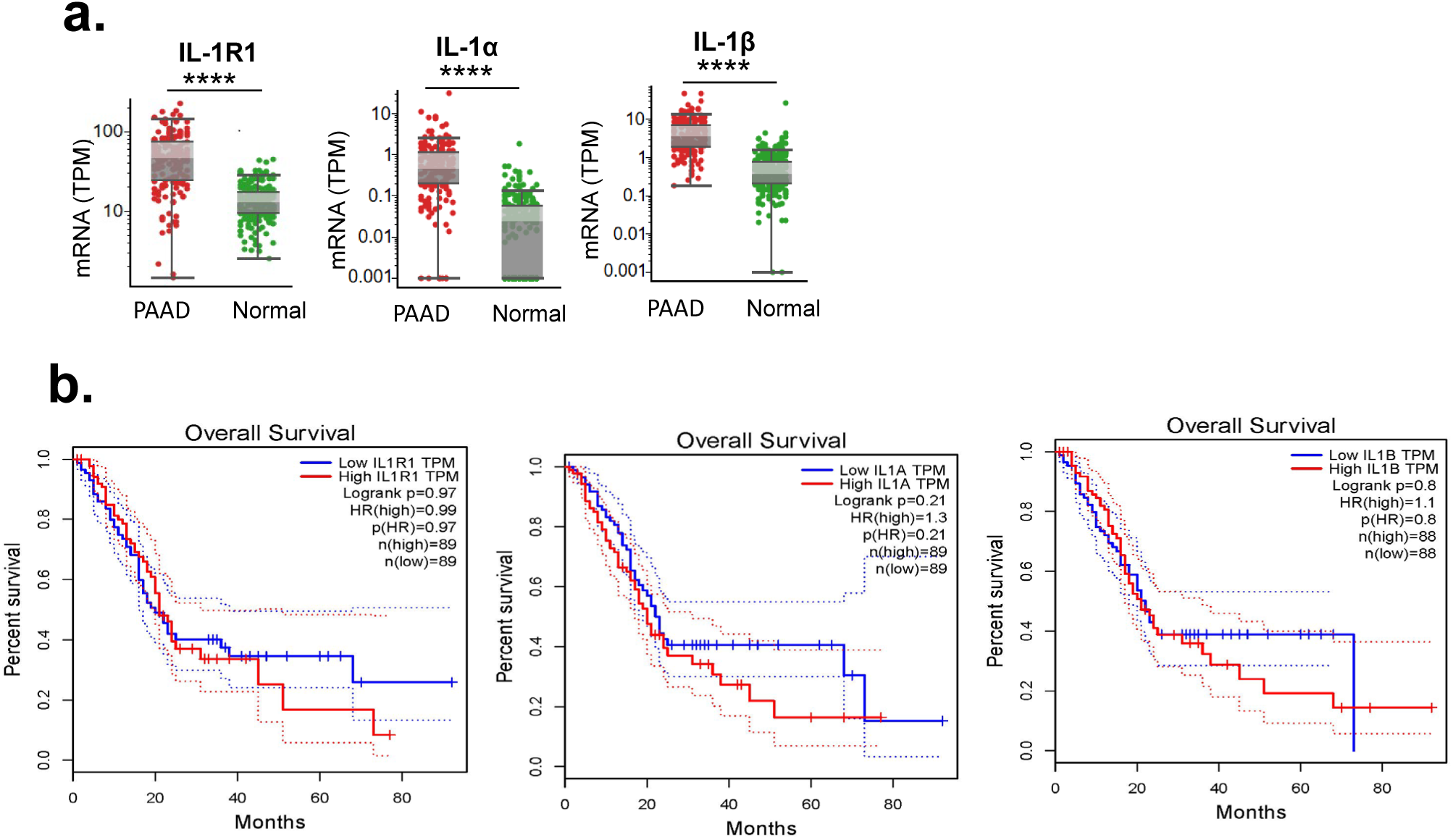
Expression of IL-1 family genes is not associated with overall survival in patients with pancreatic ductal adenocarcinoma (PDAC). (a) Box plots showing the expression levels of IL-1R1, IL-1α, and IL-1β in patient samples. Gene expression (TPM: transcripts per million) from RNA sequencing is shown in PDAC samples from The Cancer Genome Atlas (PAAD) and normal pancreatic tissue (****p < 0.0001, Student *t* test). Each dot represents an individual sample. Median expression is indicated by the horizontal line within each box, with whiskers representing the interquartile range. (b) Kaplan-Meier survival curves depicting overall survival based on high (red line) and low (blue line) expression of IL-1R1, IL-1α, and IL-1β. Dotted lines indicate 95% confidence intervals. Log-rank p values and hazard ratios (HR) for each gene are provided in the figure. IL-1R1: No significant difference in overall survival (log-rank p = 0.97, HR = 0.99). IL-1α: No significant difference in overall survival (log-rank p = 0.21, HR = 1.3). IL-1β: No significant difference in overall survival (log-rank p = 0.8, HR = 1.1). Each survival analysis was performed with nearly equal sample sizes between high and low expression groups (n ≈ 88–89 per group).

## Discussion

Agonistic CD40 antibodies have emerged as promising immunotherapeutic agents that activate both innate and adaptive antitumor immunity. Although agonistic CD40 antibodies have shown efficacy across several solid tumors, including PDAC, clinical responses remain modest, with only a few durable outcomes reported when the treatment was administered alone or in combination with immune checkpoint blockade. PMN-MDSCs (CD11b^+^Ly6C^+^Ly6G^+^ cells) are well-established contributors to the immunosuppressive tumor microenvironment in PDAC. Beyond dampening antitumor immunity, PMN-MDSCs have also been implicated in treatment-associated liver toxicity. Notably, no clinically approved therapeutics specifically target PMN-MDSCs in humans, underscoring an unmet need for strategies to mitigate their immunosuppressive effects without compromising host defenses. Based on our previous findings, we hypothesized that IL-1R1 blockade would potentiate agonistic CD40 antibody efficacy while mitigating liver toxicity in PDAC by reducing PMN-MDSC infiltration. The current study is the first to determine the effect of IL-1R1 blockade on the efficacy and toxicity of agonistic CD40 antibody therapy in PDAC.

The findings of the current study revealed striking tumor-specific differences. In contrast to melanoma, in our orthotopic PDAC models, IL-1R1 blockade failed to enhance agonistic CD40 antibody efficacy. Surprisingly, agonistic CD40 antibody monotherapy outperformed combination therapy in the subcutaneous PDAC model, and IL-1R1 blockade alone was associated with significantly worse survival compared with isotype control. These findings highlight fundamental context-dependent roles of IL-1 signaling, suggesting that its impact on tumor immunity is tumor type–specific and influenced by the local microenvironment.

Further investigations into the role of Ly6G⁺ cells (PMN-MDSCs or neutrophils) provided additional insights. Selective depletion of Ly6G⁺ cells unexpectedly compromised agonistic CD40 antibody efficacy, accelerating tumor growth and reducing survival in PDAC-bearing mice. This finding challenges the traditional view of PMN-MDSCs as purely pro-tumorigenic, suggesting that Ly6G⁺ cells may possess antitumorigenic properties within the PDAC microenvironment.

Our findings align with those of Fan et al ^31^, who showed that tumor-derived IL-1 receptor antagonist (IL-1Ra) exhibits immunosuppressive functions and promotes pancreatic cancer progression. This further underscores the complex role of IL-1 signaling in PDAC and its impact on immune modulation and tumor growth.

Recent reports showed that tumor cell–derived IL-1β induces immune suppression in the pancreatic cancer microenvironment. Das et al ^32^ reported that neutralizing IL-1β enhanced the efficacy of anti-PD-1, and Takahashi et al ^33^ reported that IL-1β promoted tumorigenesis through immune-suppressive B cells. Aggen et al recently showed that the combination of IL-1β blockade and either anti-PD-1 or a tyrosine kinase inhibitor had greater antitumor activity than either therapy alone against mouse renal cell carcinoma ^34^. We cannot directly correlate our findings with these findings because IL-1R1 blockade inhibits both IL-1α and IL-1β pathways, not solely IL-1β.

Our preclinical findings support those of Herremans et al ^35^, who reported that in PDAC data from The Cancer Genome Atlas and Gene Expression Omnibus databases, IL-1 was associated with significant changes in the PDAC tumor immune landscape and the expression of tumor immune checkpoint proteins.

Dosch et al ^36^ demonstrated that the human IL-1R1 antagonist anakinra significantly reduced IL-6 levels and inhibited STAT3 activation in pancreatic tumors derived from PKT mice. In this genetically engineered mouse model (GEMM) of PDAC, the combination of anakinra with chemotherapy markedly extended overall survival compared to either vehicle treatment or anakinra monotherapy. These findings suggest that IL-1R1 blockade can enhance chemotherapy efficacy, though its impact may be context-dependent, potentially influenced by tumor microenvironment dynamics or specific treatment regimens.

IL-1 induces liver inflammation in various liver diseases, such as alcoholic and non-alcoholic liver disease, autoimmune hepatitis, and viral hepatitis ^38^. However, the toxic effect of cancer immunotherapy-induced IL-1 on the liver, colon, and other organs has not been studied. Furthermore, although IL-6, IL-8, IL-17, and C-reactive protein have been shown to be the major cytokines that cause immune-related adverse events in response to cancer immunotherapy and are downstream of IL-1, blockade of the IL-1 pathway to suppress immunotherapy-mediated toxicity has not been explored.

Agonistic CD40 antibody therapy has been shown to induce liver damage, primarily attributed to tumor-derived PMN-MDSCs ^12,30^. The current study revealed that although IL-1R1 blockade effectively reduced PMN-MDSC levels in mice, it unexpectedly exacerbated agonistic CD40 antibody–induced hepatotoxicity. This finding suggests that CD40-induced liver toxicity is independent of IL-1 signaling or PMN-MDSCs, highlighting a more complex mechanism underlying CD40-mediated hepatotoxicity that warrants further investigation.

In conclusion, the current study underscores the complex and context-dependent nature of IL-1 signaling and PMN-MDSC function in PDAC. Although agonistic CD40 antibodies remain promising, therapeutic strategies aimed at enhancing their efficacy and lowering toxicity must consider the tumor-specific immune landscape to avoid unintended consequences.

## Materials and Methods

### Mice and tumor model

Male C57BL/6 mice (6–8 weeks old) were obtained from Jackson Laboratory and maintained under specific pathogen-free conditions at the institutional animal facility. All experiments were performed in accordance with the guidelines approved by the Institutional Animal Care and Use Committee of The University of Texas MD Anderson Cancer Center.

To establish subcutaneous PDAC tumors, we injected mice subcutaneously into the right flank with 1 × 10⁶ KPC cells (Kras^+/G12D^ TP53^+/R172H^ Pdx1-Cre), a well-established murine PDAC cell line, generously provided by Dr. Michael Curran (MD Anderson). Tumors were allowed to grow for 9 days before treatment initiation.

### Orthotopic PDAC tumor cell implantation

Orthotopic PDAC tumor implantation was performed using luciferase (Luc)-labeled KPC cells. A total of 30,000 cells were implanted following a standard protocol. Tumor growth was monitored weekly using IVIS bioluminescence imaging, as previously described ^39^.

### Treatment scheme

Tumor presence was confirmed by imaging before treatment initiation. Beginning 9 days after tumor implantation, subcutaneous and orthotopic PDAC-bearing mice were treated with anti-IL-1R1 (Clone JAMA-147, BioXCell) or isotype control antibodies (BioXCell) administered intraperitoneally. Twenty-four hours after the initial treatment, the mice received an agonistic CD40 antibody (Clone FGK4.5, BioXCell) or corresponding isotype controls administered intraperitoneally. Treatments were administered every 3 days for 2 weeks, followed by once-weekly dosing. In select experiments, mice received an anti-Ly6G antibody (depletion antibody; Clone 1A8, BioXCell) twice weekly. All antibodies were administered at a dose of 200 µg per injection intraperitoneally.

### Tumor growth and survival

Subcutaneous tumor growth was monitored using calipers, with tumor size calculated as the product of perpendicular diameters. Mice were euthanized when tumors reached 200 mm². Orthotopic tumor growth was monitored weekly using IVIS bioluminescence imaging. Mice were euthanized upon reaching a significant tumor burden. Survival analysis was performed using the log-rank test.

### Serum analysis

Blood samples were collected via retro-orbital bleeding at three time points: 48 hours after treatment initiation, after 9 days, and after 3 weeks. Serum ALT and AST levels were measured using the COBAS INTEGRA 400 Plus analyzer (Roche Diagnostics) to assess liver toxicity.

### Histological analysis

Liver and tumor tissues were harvested after 1 and 2 weeks of treatment. Tissues were fixed in 10% neutral-buffered formalin, embedded in paraffin, and sectioned. H&E staining was performed to assess histological changes, including tissue architecture, necrosis, and immune cell infiltration.

### Flow Cytometric analysis

Red blood cells lysis was performed on blood. Cells were surface stained with Abs against CD45 APC (clone104, BioLegend), CD11b FITC (clone M1/70, eBioscience), Ly6CPB (clone HK1.4, BioLegend), Ly6G PE (clone 1A8, BioLegend). Data were acquired on a Canto II flow cytometer (BD Biosciences) and analyzed using FlowJo software (version 10.10).

### Transcriptome analysis of liver and tumor

Bulk RNA sequencing was performed by Novogene (Beijing, China), a leading provider of high-throughput sequencing services. Total RNA was extracted from flash-frozen tissue samples using the RNeasy Mini Kit (Qiagen) according to the manufacturer’s instructions. RNA quality and integrity were assessed using the Agilent 2100 Bioanalyzer, ensuring RNA integrity numbers (RIN) > 7.0 for downstream analysis. Sequencing libraries were prepared using the NEBNext Ultra II RNA Library Prep Kit (New England Biolabs), incorporating poly(A) enrichment to focus on mRNA transcripts.

Libraries were quantified using Qubit and real-time PCR and sequenced on the Illumina NovaSeq 6000 platform, generating 150-bp paired-end reads. Raw data underwent quality control using Fast QC, and clean reads were aligned to the mouse reference genome (GRCm39) using HISAT2. Differential expression analysis was performed with DESeq2, and KEGG and gene ontology pathway enrichment analyses were conducted using the Cluster Profiler R package.

### Statistical analysis

Data were analyzed using GraphPad Prism 10 software. All results are expressed as mean ± SEM. Tumor growth was evaluated by two-way ANOVA with the Tukey post hoc test, and nonparametric data were analyzed using the Wilcoxon rank-sum test. Statistical significance was defined as p < 0.05.

## Data availability

The datasets generated and analyzed during the current study are available from the corresponding author upon reasonable request.

## Acknowledgments

This work was supported by the University Cancer Foundation via the Institutional Research Grant program at The University of Texas MD Anderson Cancer Center and Hirshberg Foundation for Pancreatic Cancer Research (to M.S.) and National Institutes of Health/National Cancer Institute Grant P30CA016672, which supports the flow cytometry facility at The University of Texas MD Anderson Cancer Center. We thank Erica Goodoff, Senior Scientific Editor in the Research Medical Library at The University of Texas MD Anderson Cancer Center, for editing this article.

## References

1. Mizrahi JD, Surana R, Valle JW, Shroff RT. Pancreatic cancer. Lancet. 2020;395(10242):2008–20. Epub 2020/07/01. doi: 10.1016/S0140-6736(20)30974-0. PubMed PMID: 32593337.

2. Rahib L, Smith BD, Aizenberg R, Rosenzweig AB, Fleshman JM, Matrisian LM. Projecting cancer incidence and deaths to 2030: the unexpected burden of thyroid, liver, and pancreas cancers in the United States. Cancer Res. 2014;74(11):2913–21. Epub 2014/05/21. doi: 10.1158/0008-5472.CAN-14-0155. PubMed PMID: 24840647.

3. Kabacaoglu D, Ciecielski KJ, Ruess DA, Algul H. Immune Checkpoint Inhibition for Pancreatic Ductal Adenocarcinoma: Current Limitations and Future Options. Front Immunol. 2018;9:1878. Epub 20180815. doi: 10.3389/fimmu.2018.01878. PubMed PMID: 30158932; PMCID: PMC6104627.

4. Schizas D, Charalampakis N, Kole C, Economopoulou P, Koustas E, Gkotsis E, Ziogas D, Psyrri A, Karamouzis MV. Immunotherapy for pancreatic cancer: A 2020 update. Cancer Treat Rev. 2020;86:102016. Epub 20200325. doi: 10.1016/j.ctrv.2020.102016. PubMed PMID: 32247999.

5. Vonderheide RH. CD40 Agonist Antibodies in Cancer Immunotherapy. Annu Rev Med. 2020;71:47–58. Epub 20190814. doi: 10.1146/annurev-med-062518-045435. PubMed PMID: 31412220.

6. Beatty GL, Chiorean EG, Fishman MP, Saboury B, Teitelbaum UR, Sun W, Huhn RD, Song W, Li D, Sharp LL, Torigian DA, O’Dwyer PJ, Vonderheide RH. CD40 agonists alter tumor stroma and show efficacy against pancreatic carcinoma in mice and humans. Science. 2011;331(6024):1612–6. doi: 10.1126/science.1198443. PubMed PMID: 21436454; PMCID: PMC3406187.

7. Li DK, Wang W. Characteristics and clinical trial results of agonistic anti-CD40 antibodies in the treatment of malignancies. Oncol Lett. 2020;20(5):176. Epub 2020/09/17. doi: 10.3892/ol.2020.12037. PubMed PMID: 32934743; PMCID: PMC7471753.

8. Padron LJ, Maurer DM, O’Hara MH, O’Reilly EM, Wolff RA, Wainberg ZA, Ko AH, Fisher G, Rahma O, Lyman JP, Cabanski CR, Yu JX, Pfeiffer SM, Spasic M, Xu J, Gherardini PF, Karakunnel J, Mick R, Alanio C, Byrne KT, Hollmann TJ, Moore JS, Jones DD, Tognetti M, Chen RO, Yang X, Salvador L, Wherry EJ, Dugan U, O’Donnell-Tormey J, Butterfield LH, Hubbard-Lucey VM, Ibrahim R, Fairchild J, Bucktrout S, LaVallee TM, Vonderheide RH. Sotigalimab and/or nivolumab with chemotherapy in first-line metastatic pancreatic cancer: clinical and immunologic analyses from the randomized phase 2 PRINCE trial. Nat Med. 2022;28(6):1167–77. Epub 20220603. doi: 10.1038/s41591-022-01829-9. PubMed PMID: 35662283; PMCID: PMC9205784.

9. Weiss SA, Sznol M, Shaheen M, Berciano-Guerrero MA, Couselo EM, Rodriguez-Abreu D, Boni V, Schuchter LM, Gonzalez-Cao M, Arance A, Wei W, Ganti AK, Hauke RJ, Berrocal A, Iannotti NO, Hsu FJ, Kluger HM. A Phase II Trial of the CD40 Agonistic Antibody Sotigalimab (APX005M) in Combination with Nivolumab in Subjects with Metastatic Melanoma with Confirmed Disease Progression on Anti-PD-1 Therapy. Clin Cancer Res. 2024;30(1):74–81. doi: 10.1158/1078-0432.CCR-23-0475. PubMed PMID: 37535056; PMCID: PMC10767304.

10. Johnson P, Challis R, Chowdhury F, Gao Y, Harvey M, Geldart T, Kerr P, Chan C, Smith A, Steven N, Edwards C, Ashton-Key M, Hodges E, Tutt A, Ottensmeier C, Glennie M, Williams A. Clinical and biological effects of an agonist anti-CD40 antibody: a Cancer Research UK phase I study. Clin Cancer Res. 2015;21(6):1321–8. Epub 2015/01/16. doi: 10.1158/1078-0432.CCR-14-2355. PubMed PMID: 25589626.

11. Vonderheide RH, Flaherty KT, Khalil M, Stumacher MS, Bajor DL, Hutnick NA, Sullivan P, Mahany JJ, Gallagher M, Kramer A, Green SJ, O’Dwyer PJ, Running KL, Huhn RD, Antonia SJ. Clinical activity and immune modulation in cancer patients treated with CP-870,893, a novel CD40 agonist monoclonal antibody. J Clin Oncol. 2007;25(7):876–83. Epub 2007/03/01. doi: 10.1200/JCO.2006.08.3311. PubMed PMID: 17327609.

12. Medina-Echeverz J, Ma C, Duffy AG, Eggert T, Hawk N, Kleiner DE, Korangy F, Greten TF. Systemic Agonistic Anti-CD40 Treatment of Tumor-Bearing Mice Modulates Hepatic Myeloid-Suppressive Cells and Causes Immune-Mediated Liver Damage. Cancer Immunol Res. 2015;3(5):557–66. Epub 20150130. doi: 10.1158/2326-6066.CIR-14-0182. PubMed PMID: 25637366; PMCID: PMC4420683.

13. Blake SJ, James J, Ryan FJ, Caparros-Martin J, Eden GL, Tee YC, Salamon JR, Benson SC, Tumes DJ, Sribnaia A, Stevens NE, Finnie JW, Kobayashi H, White DL, Wesselingh SL, O’Gara F, Lynn MA, Lynn DJ. The immunotoxicity, but not anti-tumor efficacy, of anti-CD40 and anti-CD137 immunotherapies is dependent on the gut microbiota. Cell Rep Med. 2021;2(12):100464. Epub 20211208. doi: 10.1016/j.xcrm.2021.100464. PubMed PMID: 35028606; PMCID: PMC8714857.

14. Hussain SP, Harris CC. Inflammation and cancer: an ancient link with novel potentials. Int J Cancer. 2007;121(11):2373–80. Epub 2007/09/26. doi: 10.1002/ijc.23173. PubMed PMID: 17893866.

15. Dinarello CA. Proinflammatory cytokines. Chest. 2000;118(2):503–8. Epub 2000/08/11. doi: 10.1378/chest.118.2.503. PubMed PMID: 10936147.

16. Voronov E, Shouval DS, Krelin Y, Cagnano E, Benharroch D, Iwakura Y, Dinarello CA, Apte RN. IL-1 is required for tumor invasiveness and angiogenesis. Proc Natl Acad Sci U S A. 2003;100(5):2645–50. Epub 2003/02/25. doi: 10.1073/pnas.0437939100. PubMed PMID: 12598651; PMCID: PMC151394.

17. Lewis AM, Varghese S, Xu H, Alexander HR. Interleukin-1 and cancer progression: the emerging role of interleukin-1 receptor antagonist as a novel therapeutic agent in cancer treatment. J Transl Med. 2006;4:48. Epub 2006/11/14. doi: 10.1186/1479-5876-4-48. PubMed PMID: 17096856; PMCID: PMC1660548.

18. Lee PY, Kumagai Y, Xu Y, Li Y, Barker T, Liu C, Sobel ES, Takeuchi O, Akira S, Satoh M, Reeves WH. IL-1alpha modulates neutrophil recruitment in chronic inflammation induced by hydrocarbon oil. J Immunol. 2011;186(3):1747–54. Epub 2010/12/31. doi: 10.4049/jimmunol.1001328. PubMed PMID: 21191074; PMCID: PMC3607541.

19. Brinster C, Shevach EM. Costimulatory effects of IL-1 on the expansion/differentiation of CD4+CD25+Foxp3+ and CD4+CD25+Foxp3-T cells. J Leukoc Biol. 2008;84(2):480–7. Epub 2008/05/15. doi: 10.1189/jlb.0208085. PubMed PMID: 18477692; PMCID: PMC2493074.

20. Cahill CM, Rogers JT. Interleukin (IL) 1beta induction of IL-6 is mediated by a novel phosphatidylinositol 3-kinase-dependent AKT/IkappaB kinase alpha pathway targeting activator protein-1. J Biol Chem. 2008;283(38):25900–12. Epub 2008/06/03. doi: 10.1074/jbc.M707692200. PubMed PMID: 18515365; PMCID: PMC2533786.

21. Jung YD, Fan F, McConkey DJ, Jean ME, Liu W, Reinmuth N, Stoeltzing O, Ahmad SA, Parikh AA, Mukaida N, Ellis LM. Role of P38 MAPK, AP-1, and NF-kappaB in interleukin-1beta-induced IL-8 expression in human vascular smooth muscle cells. Cytokine. 2002;18(4):206–13. Epub 2002/07/20. doi: 10.1006/cyto.2002.1034. PubMed PMID: 12126643.

22. Zhang D, Sun M, Samols D, Kushner I. STAT3 participates in transcriptional activation of the C-reactive protein gene by interleukin-6. J Biol Chem. 1996;271(16):9503–9. Epub 1996/04/19. doi: 10.1074/jbc.271.16.9503. PubMed PMID: 8621622.

23. Amlani A, Barber C, Fifi-Mah A, Monzon J. Successful Treatment of Cytokine Release Syndrome with IL-6 Blockade in a Patient Transitioning from Immune-Checkpoint to MEK/BRAF Inhibition: A Case Report and Review of Literature. Oncologist. 2020;25(7):e1120–e3. Epub 2020/04/28. doi: 10.1634/theoncologist.2020-0194. PubMed PMID: 32337758; PMCID: PMC7356700.

24. Yoshino K, Nakayama T, Ito A, Sato E, Kitano S. Severe colitis after PD-1 blockade with nivolumab in advanced melanoma patients: potential role of Th1-dominant immune response in immune-related adverse events: two case reports. BMC Cancer. 2019;19(1):1019. Epub 2019/10/31. doi: 10.1186/s12885-019-6138-7. PubMed PMID: 31664934; PMCID: PMC6819390.

25. Schalper KA, Carleton M, Zhou M, Chen T, Feng Y, Huang SP, Walsh AM, Baxi V, Pandya D, Baradet T, Locke D, Wu Q, Reilly TP, Phillips P, Nagineni V, Gianino N, Gu J, Zhao H, Perez-Gracia JL, Sanmamed MF, Melero I. Elevated serum interleukin-8 is associated with enhanced intratumor neutrophils and reduced clinical benefit of immune-checkpoint inhibitors. Nat Med. 2020;26(5):688–92. Epub 2020/05/15. doi: 10.1038/s41591-020-0856-x. PubMed PMID: 32405062; PMCID: PMC8127102.

26. Zhang Y, Chandra V, Riquelme Sanchez E, Dutta P, Quesada PR, Rakoski A, Zoltan M, Arora N, Baydogan S, Horne W, Burks J, Xu H, Hussain P, Wang H, Gupta S, Maitra A, Bailey JM, Moghaddam SJ, Banerjee S, Sahin I, Bhattacharya P, McAllister F. Interleukin-17-induced neutrophil extracellular traps mediate resistance to checkpoint blockade in pancreatic cancer. J Exp Med. 2020;217(12). Epub 2020/08/30. doi: 10.1084/jem.20190354. PubMed PMID: 32860704; PMCID: PMC7953739

27. Tarhini AA, Zahoor H, Lin Y, Malhotra U, Sander C, Butterfield LH, Kirkwood JM. Baseline circulating IL-17 predicts toxicity while TGF-beta1 and IL-10 are prognostic of relapse in ipilimumab neoadjuvant therapy of melanoma. J Immunother Cancer. 2015;3:39. Epub 2015/09/18. doi: 10.1186/s40425-015-0081-1. PubMed PMID: 26380086; PMCID: PMC4570556.

28. Singh S, Xiao Z, Bavisi K, Roszik J, Melendez BD, Wang Z, Cantwell MJ, Davis RE, Lizee G, Hwu P, Neelapu SS, Overwijk WW, Singh M. IL-1alpha Mediates Innate and Acquired Resistance to Immunotherapy in Melanoma. J Immunol. 2021;206(8):1966–75. Epub 20210315. doi: 10.4049/jimmunol.2000523. PubMed PMID: 33722878; PMCID: PMC8023145.

29. Jaillon S, Ponzetta A, Di Mitri D, Santoni A, Bonecchi R, Mantovani A. Neutrophil diversity and plasticity in tumour progression and therapy. Nat Rev Cancer. 2020;20(9):485–503. Epub 20200721. doi: 10.1038/s41568-020-0281-y. PubMed PMID: 32694624.

30. Kapanadze T, Medina-Echeverz J, Gamrekelashvili J, Weiss JM, Wiltrout RH, Kapoor V, Hawk N, Terabe M, Berzofsky JA, Manns MP, Wang E, Marincola FM, Korangy F, Greten TF. Tumor-induced CD11b(+) Gr-1(+) myeloid-derived suppressor cells exacerbate immune-mediated hepatitis in mice in a CD40-dependent manner. Eur J Immunol. 2015;45(4):1148–58. Epub 20150223. doi: 10.1002/eji.201445093. PubMed PMID: 25616156; PMCID: PMC4425346.

31. Fan YC, Fong YC, Kuo CT, Li CW, Chen WY, Lin JD, Burtin F, Linnebacher M, Bui QT, Lee KD, Tsai YC. Tumor-derived interleukin-1 receptor antagonist exhibits immunosuppressive functions and promotes pancreatic cancer. Cell Biosci. 2023;13(1):147. Epub 20230810. doi: 10.1186/s13578-023-01090-8. PubMed PMID: 37563620; PMCID: PMC10416534.

32. Das S, Shapiro B, Vucic EA, Vogt S, Bar-Sagi D. Tumor Cell-Derived IL1beta Promotes Desmoplasia and Immune Suppression in Pancreatic Cancer. Cancer Res. 2020;80(5):1088–101. Epub 20200108. doi: 10.1158/0008-5472.CAN-19-2080. PubMed PMID: 31915130; PMCID: PMC7302116.

33. Takahashi R, Macchini M, Sunagawa M, Jiang Z, Tanaka T, Valenti G, Renz BW, White RA, Hayakawa Y, Westphalen CB, Tailor Y, Iuga AC, Gonda TA, Genkinger J, Olive KP, Wang TC. Interleukin-1beta-induced pancreatitis promotes pancreatic ductal adenocarcinoma via B lymphocyte-mediated immune suppression. Gut. 2021;70(2):330–41. Epub 20200510. doi: 10.1136/gutjnl-2019-319912. PubMed PMID: 32393543.

34. Aggen DH, Ager CR, Obradovic AZ, Chowdhury N, Ghasemzadeh A, Mao W, Chaimowitz MG, Lopez-Bujanda ZA, Spina CS, Hawley JE, Dallos MC, Zhang C, Wang V, Li H, Guo XV, Drake CG. Blocking IL1 Beta Promotes Tumor Regression and Remodeling of the Myeloid Compartment in a Renal Cell Carcinoma Model: Multidimensional Analyses. Clin Cancer Res. 2021;27(2):608–21. Epub 20201104. doi: 10.1158/1078-0432.CCR-20-1610. PubMed PMID: 33148676; PMCID: PMC7980495.

35. Herremans KM, Szymkiewicz DD, Riner AN, Bohan RP, Tushoski GW, Davidson AM, Lou X, Leong MC, Dean BD, Gerber M, Underwood PW, Han S, Hughes SJ. The interleukin-1 axis and the tumor immune microenvironment in pancreatic ductal adenocarcinoma. Neoplasia. 2022;28:100789. Epub 20220405. doi: 10.1016/j.neo.2022.100789. PubMed PMID: 35395492; PMCID: PMC8990176.

36. Dosch AR, Singh S, Dai X, Mehra S, Silva IC, Bianchi A, Srinivasan S, Gao Z, Ban Y, Chen X, Banerjee S, Nagathihalli NS, Datta J, Merchant NB. Targeting Tumor-Stromal IL6/STAT3 Signaling through IL1 Receptor Inhibition in Pancreatic Cancer. Mol Cancer Ther. 2021;20(11):2280–90. Epub 20210913. doi: 10.1158/1535-7163.MCT-21-0083. PubMed PMID: 34518296; PMCID: PMC8571047.

37. Theivanthiran B, Evans KS, DeVito NC, Plebanek M, Sturdivant M, Wachsmuth LP, Salama AK, Kang Y, Hsu D, Balko JM, Johnson DB, Starr M, Nixon AB, Holtzhausen A, Hanks BA. A tumor-intrinsic PD-L1/NLRP3 inflammasome signaling pathway drives resistance to anti-PD-1 immunotherapy. J Clin Invest. 2020;130(5):2570–86. doi: 10.1172/JCI133055. PubMed PMID: 32017708; PMCID: PMC7190922.

38. Barbier L, Ferhat M, Salame E, Robin A, Herbelin A, Gombert JM, Silvain C, Barbarin A. Interleukin-1 Family Cytokines: Keystones in Liver Inflammatory Diseases. Front Immunol. 2019;10:2014. Epub 2019/09/12. doi: 10.3389/fimmu.2019.02014. PubMed PMID: 31507607; PMCID: PMC6718562.

39. Ager CR, Boda A, Rajapakshe K, Lea ST, Di Francesco ME, Jayaprakash P, Slay RB, Morrow B, Prasad R, Dean MA, Duffy CR, Coarfa C, Jones P, Curran MA. High potency STING agonists engage unique myeloid pathways to reverse pancreatic cancer immune privilege. J Immunother Cancer. 2021;9(8). Epub 2021/08/04. doi: 10.1136/jitc-2021-003246. PubMed PMID: 34341132; PMCID: PMC8330562.

